# Site-specific immunogenicity and anti-PD1 response in mismatch repair deficient lung adenocarcinoma models

**DOI:** 10.1101/2025.01.13.632715

**Authors:** Etna Abad, Ivan Zadra, Anastasia Krasko, Marc Subirana-Granés, Diana Reyes, Pablo Borredat, Lorenzo Pasquali, Jose Aramburu, Cristina López-Rodríguez, Ana Janic

**Affiliations:** Department of Medicine and Life Sciences, Pompeu Fabra University, Barcelona, Spain

## Abstract

Immunotherapy has revolutionized cancer treatment, yet responses vary significantly based on tumour characteristics and microenvironment. Here, we developed and analysed subcutaneous and orthotopic immunocompetent mice models of mismatch repair-deficient (dMMR) lung adenocarcinoma (LUAD) by selectively ablating Mlh1. Subcutaneous tumours demonstrated partial sensitivity to anti-PD1 therapy, characterized by tumour volume reduction without significant changes in immune infiltration. In contrast, orthotopic tumours exhibited robust responses, with substantial reductions in tumour burden, enhanced immune infiltration, and increased CD4^+^ memory T cells, highlighting the critical role of anatomical site and tumour microenvironment in shaping immunotherapy outcomes. Our findings emphasise the relevance of orthotopic models for preclinical evaluation and suggest that they more accurately reflect clinical responses to immune checkpoint blockade in dMMR LUAD.

## Introduction

Immunotherapy has emerged as a revolutionary approach in the treatment of various malignancies. In recent years, immune checkpoint inhibitors have been demonstrating significant efficacy across multiple cancer types[1,2]. Still, the response to immunotherapy is not uniform across all tumours, leading to the classification of “hot” and “cold” tumours based on their immunogenicity and susceptibility to immune-mediated elimination[3]. Recent research has exposed a correlation between genomic instability and immunotherapy response. Tumours with high levels of genomic instability often have a higher tumour mutational burden (TMB), leading to the presentation of more neoantigens and triggering a stronger immune response[4]. Consequently, high TMB has been recently accepted as a potential biomarker for identifying patients who may benefit most from anti-PD1 treatment[5,6].

Among genomically unstable tumours, those with mismatch repair deficiency (dMMR) in which microsatellite instability (MSI) often occurs throughout the genome, have demonstrated notable positive responses to immunotherapy[7]. The success of immunotherapy in dMMR tumours has led to FDA approvals for certain immune checkpoint inhibitors (ICB) across multiple cancer types. Specifically, anti-PD1 therapy has received approval for all dMMR/MSI-high solid tumours, regardless of the tissue of origin[8].

However, it is important to note that even among dMMR tumours, not all patients respond to immunotherapy[7,9]. Several factors may contribute, among them, the tumour microenvironment, the presence of other immune-suppressive mechanisms, the degree of T-cell infiltration or the mutational heterogeneity of the tumour populations, giving rise to different concentrations of neoantigens exposed to the immune system[10].

There is a need to improve the preclinical immunocompetent models for evaluating immunotherapy efficacy, as it is associated with unique factors like specific immune infiltration. To address this, we developed orthotopic and subcutaneous immunocompetent models of dMMR lung adenocarcinoma (LUAD) tumours by selectively ablating MLH1 in LUAD murine cells. Our findings indicate that implantation site differences significantly influence preclinical modelling of immune checkpoint inhibitor therapies and suggest that the immunogenic orthotopic model may more accurately reflect clinical responses to anti-PD1 treatment in dMMR lung adenocarcinoma.

## Results

### Sensitivity of Mlh1 deficient LUAD subcutaneous tumours to PD1 antibody

In this study we developed a murine tumour model in which we have knocked out Mlh1, whose inactivation contributes to MSI in cancers [12] and is widely used by the scientific community since it demonstrates sensitivity to anti-PD1[25]. Starting with commonly used poorly immunogenic *KRAS^G12D^* mutant and *TRP53* knockout lung cancer cells (KP), derived from an autochthonous mouse model of *Kras^LSL-G12D^; Trp53^flox/flox^* mice [11] we knocked down Mlh1 by shRNA (hereafter shMlh1^KP^). The Mlh1 knockdown was confirmed by Western blot analysis (Supplementary Figure 1A).

First, we compared sensitivity to anti-PD1 treatment of tumours developed by subcutaneous implantation of GFP labeled shMlh1^KP^ cells into syngeneic immunocompetent mice (C57BL/6). We chose 3 times a week intraperitoneal administration of 100µg anti-PD1 antibody or IgG2a isotope control (Figure 1A). As expected, we obtained a reduction in tumour size upon anti-PD1 treatment compared to control animals (Figure 1B-D). Surprisingly, despite the overall reduction in tumour volume in anti-PD1 *vs* control treated mice, we found that the proportion of cancer cells remained relatively unchanged (Figure 1E). Similarly, our analysis of tumour infiltrating lymphocytes revealed no substantial changes in the percentages of CD45^+^ cells, including CD4^+^ and CD8^+^ T cells or their subpopulations (effector, memory, naïve) following the treatment (Figure 1F-J, Supplementary Figure 1B,C). Likewise, no differences were detected with regards to the proportion of tumour-associated macrophages (TAMs), including TAM A, B or C (Supplementary Figure 1D) nor in dendritic cells infiltration (cDC1 or cDC2) (Supplementary Figure 1E).

**Figure 1.**
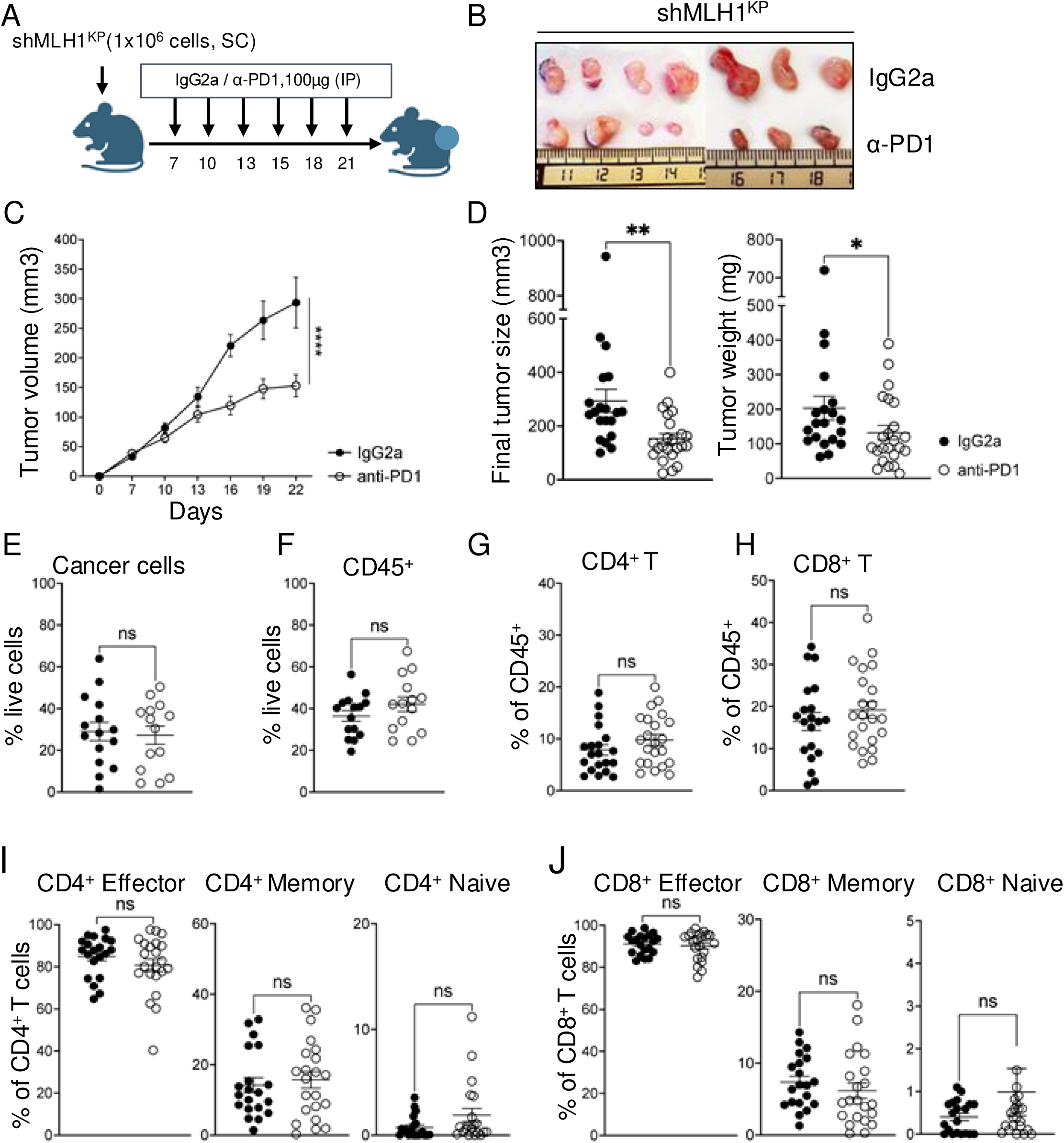
Sensitivity of Mlh1 deficient LUAD subcutaneous tumours to PD1 antibody. (A) Schedules for anti-PD1 treatment. (B) Representative subcutaneous tumour images at ethical endpoint. (C) Growth of subcutaneous shMLH1^KP^ tumours treated with anti-PD1 and IgG2a. N=20 for IgG2a, N=23 anti-PD treated mice. ****P ≤ 0.0001, two-way ANOVA. (D) Final measurement of tumour volume (left) and weight (right) at ethical endpoint. N=20 for IgG2a, N=23 anti-PD1 treated mice (from 3 independent experiments). **P≤ 0.01, *P≤ 0.05, Mann Whitney test. (E) Percentage of GFP^+^ cancer cells (N=15 IgG2a, N=14 anti-PD1), (F) CD45^+^ (N=15 IgG2a, N=14 anti-PD1), (G) CD4^+^ T cells and (H) CD8^+^ T cells (N=20 IgG2a,N=22 anti-PD1), (J) effector memory (effector), central memory (memory) or naïve cells in the total CD4^+^ T or CD8^+^T population (N=20 IgG2a, N=22 anti-PD1) analysed by flow cytometry. n.s= non-significant, **P≤ 0.01, *P≤ 0.05, two-tailed unpaired t-test or Mann Whitney depending on data normality test. Data is represented as Mean ± SEM.

### Increased tumour mutation burden and genomic alterations in Mlh1 deficient LUAD tumours treated with anti-PD1 therapy

To confirm MSI phenotype and investigate the degree of tumour mutation burden (TMB), we performed whole genome sequencing (WGS) on dissected subcutaneous tumours, including subcutaneous shMlh1^KP^ tumours that have been treated with anti-PD1 or with isotype control. We also performed WGS on shMlh1^KP^ and Mlh1 proficient WT^KP^ cell line grown *in vitro*. shMlh1^KP^ cells *in vivo* and *in vitro* showed increased burden of INDELs, with most common MSI mutational signatures ID1, ID2 and ID12[26] (Figure 2A). The number of unique variants found was increased in the shMlh1^KP^ cell line (n°=183.208) compared with the original WT^KP^ (n°=150.466) and it was unchanged upon IgG2a control treatment (average of n°=178.924 variants) (Figure 2B). Interestingly, anti-PD1 treated shMlh1^KP^ tumours have increased presence of unique variants (average of n°=255.376), mostly due to increased number of INDELs (Figure 2B). These changes were also replicated when evaluating the TMB, which was increased in anti-PD1 treated tumours (TMB average fold increase 26,36) (Figure 2C). Among clonal mutations detected in the post-treatment tumours included recurrent nonsynonymous mutation in *Ush2a* and *Nlrp1b* genes consistent with previous observations that mutations in *Ush2a* and *Nlrp1b* are implicated in signalling pathways involved in the immune system and cancer progression[27–29] (Figure 2D).

**Figure 2.**
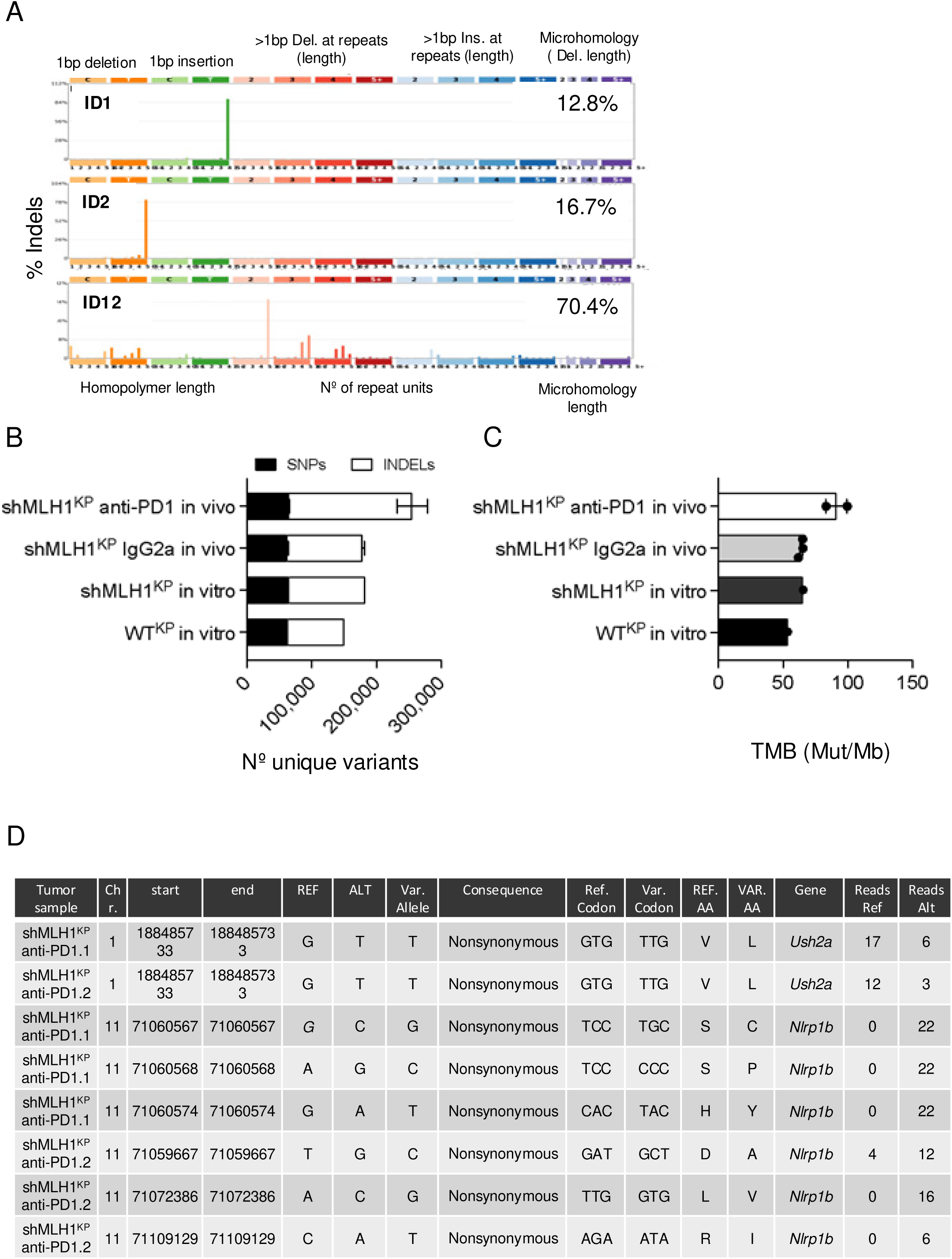
**Increased tumour mutation burden and genomic alterations in Mlh1 deficient LUAD tumours treated with anti-PD1 therapy** (A) ID mutational signatures observed in shMLH1^KP^ subcutaneous tumours are decomposed and matched to COSMIC annotations. The ID mutation signature profile of shMLH1^KP^ subcutaneous tumours corresponds to those associated with DNA mismatch repair and microsatellite instability (MSI), specifically ID1 and ID2. (B) Number of unique variants found in samples from *in vitro* WT^KP^ (N=1) or shMLH1^KP^ cell lines (N=1) and from cells recovered from tissue samples after receiving either IgG2a or anti-PD1 treatment (N=3 for IgG2a, N=2 for anti-PD1 treated tumours). Variants are shown as SNPs or INDELs. (C) TMB (Mut/Mb) of *in vitro* WT^KP^ or shMLH1^KP^ cell lines (N=1) and from cells recovered from tissue samples after receiving either IgG2a or anti-PD1 treatment (N=3 for IgG2a, N=2 for anti-PD1 treated tumours. (D) Table showing unique variants found only in cells recovered from anti-PD1 treated samples, compared with IgG2a-treated controls. Table shows sample Id, chromosomal location, reference nucleotide, altered nucleotide, consequences of mutation, reference codon, variant codon, reference amino acid, variant amino acid, gene affected, and number of reads pertaining to reference or to alternative variants.

These results highlight the effectiveness of our subcutaneous shMlh1^KP^ model in replicating the key mutational processes driving hypermutation in dMMR human cancers. It exhibits a partial response to anti-PD1 therapy, as evidenced by an overall reduction in tumour volume, but without major shifts in the cellular balance of cancer cells and immune cell infiltrates when compared to non-treated controls.

### Sensitivity of Mlh1 deficient LUAD orthotopic tumours to PD1 antibody

We hypothesised that the moderate immunogenicity in shMlh1^KP^ subcutaneous tumours might be due to the subclonality of mutations and the implantation site, both of which have been shown to be less efficient at generating effective adaptive immune responses [30–33]. First, we established clonal cell lines from the KP bulk cell line, which were subsequently single-cell-cloned to increase the frequency of clonal neoantigens. The immunogenicity of three different single cell KP clones was assessed by comparing the growth of cells subcutaneously transplanted into syngeneic immunocompetent (C57BL/6) and immune-deficient (NSG) mice (Supplementary Figure 2A). All three KP clones tested grew significantly slower in C57BL/6 compared to NSG mice. We selected one of the evaluated clones for further experiments. Using CRISPR technology, we edited the selected KP clonal cell line to generate Mlh1 deficiency (hereafter sgMlh1^KP^). Immunoblot and qRT-PCR analysis showed that Mlh1 was successfully knocked out (Supplementary Figure 2B,C).

To assess whether the sensitivity to PD1 antibody was dependent on the anatomic site of tumour growth and subclonality of the cell line, we treated orthotopic sgMlh1^KP^ lung tumours with anti-PD1. GFP labelled sgMlh1^KP^ cells were injected intravenously into mice, which were subsequently treated 7 days after engraftment (Figure 3A). Similarly to subcutaneous tumours, orthotopic lung tumours responded to anti-PD1 therapy, resulting in a significant decrease in lung tumour growth compared to control tumour-bearing mice (Figure 3B-D). Importantly, we found that the percentage of cancer cells was significantly reduced at the endpoint in anti-PD1 treated mice, (Figure 3E) which contrasts with the subcutaneous lung tumours (Figure 1E) and indicates a more robust anti-tumour response in the orthotopic setting.

**Figure 3.**
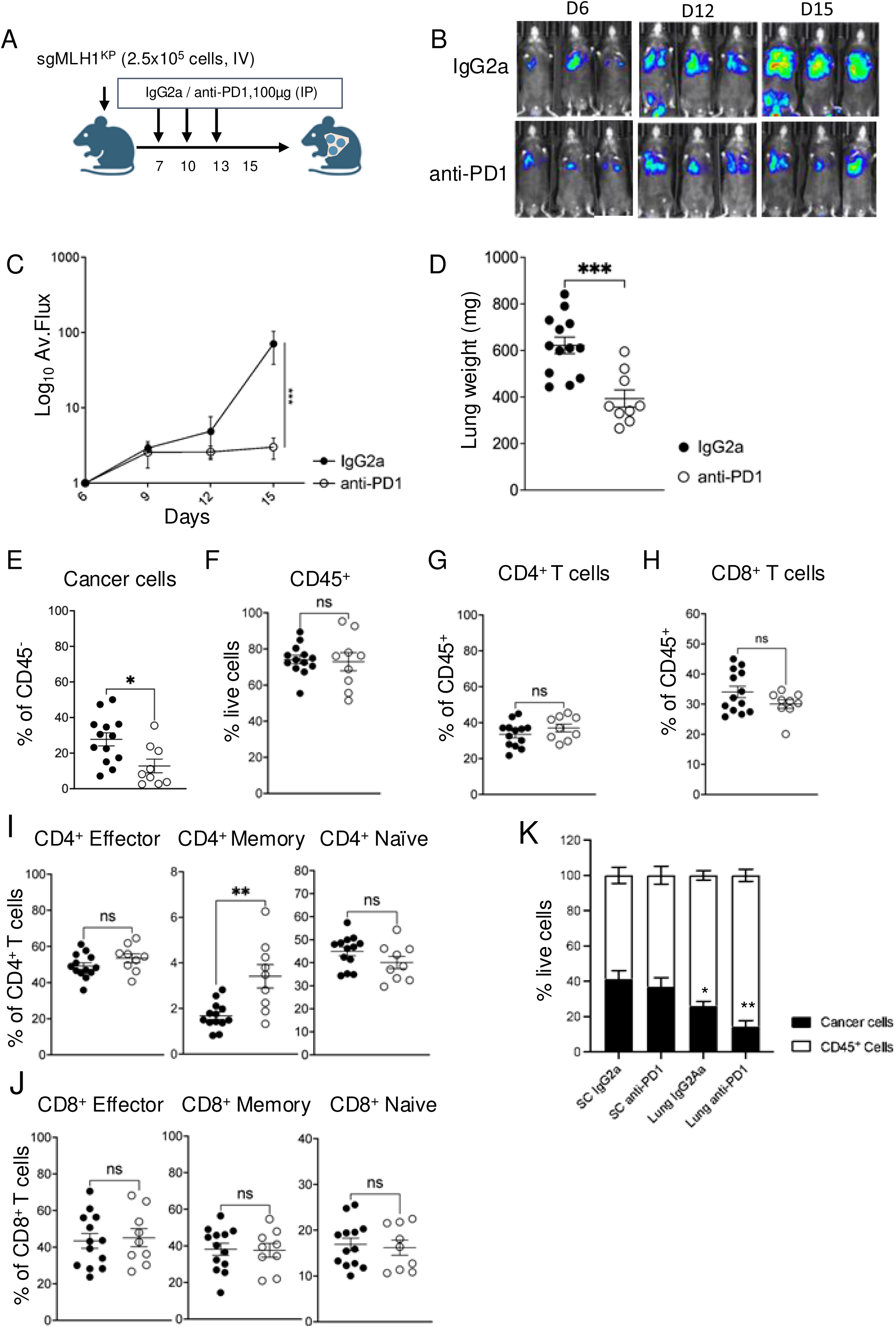
**Sensitivity of Mlh1 deficient LUAD orthotopic tumours to PD1 antibody** (A) Schedules for anti-PD1 treatment. (B) IVIS images of animals with tumour burden located in the lungs. (C) Average IVIS radiance (Log10). N=4 IgG2a, N=4 anti-PD1 treated mice. ***P ≤ 0.001, two-way ANOVA. (D) Measurements of lung weight at the experimental endpoint. N=13 IgG2a, N=9 anti-PD1 treated mice. (E-J) Flow cytometry analysis from mice treated with anti-PD1 and IgG2a control showing the percentage of GFP^+^ cancer cells, CD45^+^, CD4^+^ T cells or CD8^+^ T cells in the total CD45^+^ cell population, and effector memory (effector), central memory (memory) or naïve cells in the total CD4^+^ T or CD8^+^ T population, respectively. N=13 IgG2a, N=9 anti-PD1 treated mice. **P≤ 0.01, *P≤ 0.05, two-tailed unpaired t-test. (K) Comparison of the proportion of cancer cells and CD45^+^ cells in reference to live cells of subcutaneous and orthotopic Mlh1 deficient LUAD models, IgG2a or anti-PD1-treated. Values are normalized to the sum of cancer and CD45^+^ cells. N=15 for IgG2a-treated subcutaneous tumours, N= 14 for anti-PD1-treated subcutaneous tumours, N= 13 for orthotopic IgG2a-treated tumours and N=9 for orthotopic anti-PD1-treated mice. n.s= non-significant,**P≤ 0.01, *P≤ 0.05, two-way ANOVA. Data is represented as Mean ± SEM.

The overall immune infiltrate, as measured by CD45^+^ cells, remained unchanged between treatment groups in lungs (Figure 3F). Analysis of T cell populations revealed similarities to our subcutaneous model findings. We did not observe significant differences in the overall percentages of CD4^+^ or CD8^+^ T cells between control and anti-PD1 treated mice (Figure 3G-J). However, we noted a significant increase in CD4^+^ Memory T cells in the anti-PD1 treated group (Figure 3I), suggesting a potential enhancement of the adaptive immune response. Number of TAMs and dendritic cells in the lungs remained without significant differences (Supplementary Figure 2D,E).

Overall, in the orthotopic model, lung tissues exhibited a higher proportion of immune cells (∼80% CD45^+^) compared to the subcutaneous model (∼60%), reflecting the lung’s more robust immune landscape. Furthermore, the proportion of CD45^+^ cells increased upon anti-PD1 treatment only in the orthotopic LUAD model (Figure 3K).

In summary, we have shown that the orthotopic model of dMMR lung adenocarcinoma demonstrated an improved response to anti-PD1 therapy compared to the subcutaneous model. We observed a significant reduction in tumour burden in the lungs followed by decrease in the percentage of cancer cells. These findings highlight that the anti-PD1 response was dependent on the site of tumour growth and suggest that the immunogenic orthotopic model may more accurately reflect the potential clinical response to anti-PD1 treatment in dMMR lung adenocarcinoma.

## Discussion

In this article we report the significant influence of tumour implantation site and subclonality on the immunogenicity and therapeutic response of dMMR lung adenocarcinoma models to anti-PD1 treatment. By establishing and analysing both subcutaneous and orthotopic murine models of dMMR LUAD, we have elucidated several critical factors that modulate the efficacy of anti-PD1 therapies.

The subcutaneous implantation of shMlh1^KP^ tumours demonstrated partial sensitivity to anti-PD1 therapy, evidenced by a reduction in overall tumour volume. However, the lack of substantial changes in immune cell infiltration or viable cancer cell proportions post-treatment highlights limitations of this model in replicating clinical outcomes. While subcutaneous tumours exhibited an increased tumour mutational burden, a recognized biomarker of immune checkpoint blockade response, this did not translate into robust anti-tumour immunity. These results suggest that despite the mutational landscape favouring immunogenicity, the tumour microenvironment in subcutaneous settings may not adequately support effective immune activation. Factors such as lower immune cell infiltration and reduced clonal neoantigen presentation likely contributed to the suboptimal response observed.

In contrast, orthotopic sgMlh1^KP^ tumours presented a markedly improved response to anti-PD1 therapy, with significant reductions in tumour burden and cancer cell population. This enhanced efficacy correlates with the robust immune microenvironment in the lung of untreated mice, characterized by higher overall immune cell infiltration, increased abundance of CD4^+^ and CD8^+^ T cells and a balanced distribution of effector memory and naïve T cells compared with the subcutaneous model. Mlh1 deficient orthotopic LUAD model also revealed a greater proportion of TAMs compared with the subcutaneous model, particularly TAM A, which are associated with improved immune responses[34]. The ability of the dMMR LUAD orthotopic model to elicit a more clinically relevant immune response underscores the importance of anatomical context in preclinical evaluations of anti-PD1 therapies.

The observed differences between subcutaneous and orthotopic models align with previous studies demonstrating the critical role of the TME and anatomical site in shaping immune responses to cancer therapies[35]. Notably, in the untreated dMMR LUAD orthotopic model, the enhanced immune infiltration, particularly of T cells and TAMs, highlights the potential for leveraging the natural immune landscape of the lung to achieve better therapeutic outcomes. Additionally, the increase in CD4^+^ memory T cells in the orthotopic model following anti-PD1 therapy suggests a more robust and sustained adaptive immune response, further supporting its translational relevance.

Our results also emphasize the need to consider clonal heterogeneity when modelling dMMR cancers[30]. The improved immunogenicity and response in orthotopic models were achieved using clonal cell lines with increased neoantigen frequency. In summary, this study highlights the critical impact of tumour implantation site and subclonal composition on the efficacy of anti-PD1 therapy in dMMR LUAD model. Our findings demonstrate orthotopic models as a standard in pre-clinical immunotherapy research, ultimately improving therapeutic strategies for dMMR cancers.

## Methods

### Cell culture

Murine lung adenocarcinoma cells *KRAS^G12D^* mutant and *TRP53* knockout (KP) cells were derived from *Kras^LSL-G12D^; Trp53^flox/flox^* mice[11] (gift from Dr. Kate Sutherland, WEHI, Australia). Cells were grown in DMEM-F12 Glutamax (12634028, Thermofisher, Spain), containing 10% Fetal Bovine Serum (FBS, S181BH, Biowest, France), 100 µg/ml penicillin/ streptomycin (15140122, Thermofisher), 4 µg /ml Hydrocortisone (H0888, Merck, Spain) and Insulin-Transferrin-Selenium mix 1x (15383661, Thermofisher). HEK293T cells were grown in DMEM (L0102, Biowest) with 10% FBS and 100 µg/ml penicillin/streptomycin. Cells were grown in 5% CO2 at 37°C. All cell lines were regularly tested for mycoplasma. Only mycoplasma-negative cells were used.

### Virus production and transduction

To generate KP cells deficient for MLH1, cells were infected with retroviral vectors containing an shRNA targeting MLH1 (shMlh1^KP^). The GFP-labeled shRNA’s is as follows: TGCTGTTGACAGTGAGCGCCAGGCATTAGTTTCTCAGTTATAGTGAAGCCACAGATGTATAACTGAGAAACTAATGCCTGATGCCTACTGCCTCGGA. shRNAs were generated into LMS (LTR/MCSV/SV40-puro-IRES-GFP) retroviral vector[12]. The viral supernatants were produced in HEK293T cells by calcium phosphate precipitation, as described in[13]. KP cells were infected, and after 48-72h, cells were FACS-sorted for GFP fluorescence using a BD Influx cell sorter (BD Biosciences, Switzerland).

To generate the isogenic MLH1-KO clone (sgMlh1^KP^), the transient plasmid PX330-iRFP[14] carrying Cas9 and a sgRNA against MLH1 (5’-CAGTTTAAGCACCGCTGTGA-3’) was used. Briefly, cells were transfected using Lipofectamine 2000 (11668030, Thermofisher) following the manufacturer’s protocol. 48h after, cells positive in iRFP fluorescence were sorted and seeded as isogenic clones. MLH1 knockout was confirmed by Sanger sequencing and downregulation of transcript and protein by RT-qPCR and Western blot, respectively. To generate cells expressing luciferase-GFP, we used the lentiviral vector pLEX-hFL2iG (gift from Dr. Antoni Celià-Terrassa, Hospital Research Mar) and sorted for GFP positive cells.

### qRT-PCR

mRNA was isolated with TRIzol Reagent (15596018, Thermofisher). Total mRNA was quantified with a NanoDrop spectrophotometer and retro-transcribed into cDNA with the Superscript IV (18090010, Thermofisher). cDNA was used for quantitative PCR with reverse transcription (RT-qPCR) analysis using SYBR green (4707516001, Thermofisher). PCR conditions were as follows: 30s at 95 °C, 40 cycles of 10s at 95 °C, 30s at 60 °C, and 5s at 72 °C. The primers for MLH1 were as follows: Forward 5’-GGGAGGACTCTGATGTGGAA-3’, Reverse 5’-AGAGCTTGGTCTGGTGCTGT-3’. The primers for housekeeping ß-actin are the following: Forward 5’-AGACTTCGAGCAGGAGATGG-3 and Reverse 5‘AGGTCTTTACGGATGTCAACG-3’.

### Western Blot

Western blot was performed as in[13]. The following primary antibodies were used: mouse anti-Mlh1 (sc-271978, SCBT, Germany), diluted 1:1000, and mouse ß-actin (sc-47778, SCBT), diluted 1:3000, The secondary antibody anti-mouse (sc-516102, SCBT) was used.

### Animal experiments

All animal experiments are compliant with ethical regulations regarding animal research and were conducted under the approval of the Ethics Committee for Animal Experiments (CEEA-PRBB, Barcelona, Spain). Subcutaneous tumour models were performed by injection of 1×10^6^ of shMLH1^KP^ cells suspended in 100 μl of PBS of 7–10-week-old male C57B6/J mice (Charles River). Tumours were grown for approximately 3 weeks and harvested at the endpoint. Treatment with either anti-PD1 (P372, Leinco Tech., Missouri, United States) or IgG2 (R1367, Leinco Tech.) (100 µg per mouse) was initiated 1 week after cell injection and administered every 72h via intraperitoneal injection. This procedure was performed in 3 independent experiments. Subcutaneous tumour growth was followed by calliper measurements and the following formula applied to measure tumour volume: volume = 1/2(length × width^2^). In the case that tumours did not grow in the flank, measurement was excluded from the comparative analysis. All animals were euthanized before or at the moment of achieving maximum tumour volume.

For the lung orthotopic murine models, 100 µl of PBS containing 250.000 sgMLH1^KP^ cells expressing Luciferase-GFP were injected in the tail vein of the mice and animals were followed by IVIS luminescence until the end of the experiment. Treatment with either anti-PD1 or IgG2 (100 µg per mouse) was initiated 1 week after cell injection and administered every 72h. Animal weight and health was controlled during all the experiment.

### Tissue processing

At the end of the experiment, subcutaneous tumours or lungs were harvested, weighted, minced and placed into either 50 ml tube containing 2.5 ml DMEM with 0.5 mg/ml of collagenase A (10103578001, Roche, Spain) and 0.01% DNase I (D4263, Merck), for subcutaneous tumours, or into gentleMACS C tubes (Miltenyi Biotec, 130-093-237) containing 3 ml of PBS 1x, 5% FBS, and 2 mg/ml collagenase A type IV (Sigma-Aldrich, C4-22-1G) for lung tissues. The lungs were carefully minced using the gentleMACS Tissue Dissociator (130-096-427, Miltenyi Biotec, Spain), and both subcutaneous tumours or lungs were digested for 40 min at 37°C under continuous rotation and then filtered through 70 µm cell strainers. The filter was then washed by adding 5 ml of complete DMEM twice, and the digested tumours were incubated with 0.5-1 ml of RBC lysis buffer (11814389001, Merck) for 2 min at 4°C, then washed with 5 ml PBS containing 10% FBS and 0.1% sodium azide (PSA buffer). Tumour pellets were resuspended in PSA to obtain single-cell tumour suspensions to proceed with flow cytometry staining or sorting as in[15].

### Flow cytometry

Single cell suspensions were resuspended in PSA and incubated for 30 min at 4 °C with 0.5 µg/1×10^6^ cells of anti-CD16/32 antibody. After washing with PSA, cells were incubated with surface marker-specific antibodies (0.2-1 μg of antibody per 1×10^6^ cells, upon titration) for 30 min in the dark at 4 °C. After staining, cells were washed with PSA. The following antibodies were used:

| Antibody | Fluorochrome | Brand | Reference | Clone |
| --- | --- | --- | --- | --- |
| CD103 | Pe-Cy7 | Biolegend | 121426 | 2E7 |
| CD11b | BV605 | Biolegend | 101257 | M1/70 |
| CD11c | PE/Dazzle 594 | Biolegend | 117347 | N418 |
| CD4 | PE-Cy5 | eBioscience | 15-0042-83 | RM4-5 |
| CD44 | APC-Cy7 | Biolegend | 103028 | IM7 |
| CD45.2 | PerCP-Cy5.5 | Biolegend | 109828 | 104 |
| CD45.2 | PE/Dazzle 594 | BioLegend | 109846 | 104 |
| CD62L | APC | eBioscience | 17-0621-83 | MEL-14 |
| CD8 | PE | Biolegend | 100708 | 53-6.7 |
| CD90.2 | Pe-Cy7 | Biolegend | 140310 | 53-2.1 |
| F4/80 | APC-eFluor780 | eBioscience | 47-4801-82 | BM8 |
| Ly6C | PE | eBioscience | 12-5932-82 | HK1.4 |
| Ly6G | FITC | Biolegend | 11-9668-82 | 1A8 |
| MHCII | APC | eBioscience | 17-5321-82 | M5/114.15.2 |
| PD-1 | BV605 | Biolegend | 135220 | 29F.1A12 |
| CD103 | Pe-Cy7 | Biolegend | 121426 | 2E7 |
| CD11b | BV605 | Biolegend | 101257 | M1/70 |

Flow cytometry analysis and fluorescence-activated cell sorting (FACS) were performed with an LSR FORTESSA equipped with a 488 nm laser, a 561 nm laser, a 633 nm laser and 405 nm laser. For cell sorting, samples were filtered through a 75 µm nylon mesh (352235, Thermofisher), the BD FACSARIA cell sorter equipped with 488 nm, 561 nm, 633 nm and 405 nm lasers. Data analysis was done using the FACSDiva version 6.2 software (BD biosciences) and SpectroFlo version 3.3.0 (Cytek Biosciences). The gating strategy can be found in Supplementary Figure 1.

### Whole Genome Sequencing

Sorting of cancer cells from subcutaneous tumours was performed by selecting GFP^+^/CD45^-^ cells in the BD FACSARIA cell sorter. Between 40.000 and 160.000 cells were recovered, depending on the sample and treatment received. Cells were lysed and DNA extracted with the QIAamp DNA Micro Kit (56304, Qiagen, Germany). Purified DNA was sent to Novogene (UK) for Whole genome sequencing analysis (NovaSeq X Plus PE150).

### Alignment and SNV/INDEL inference

FASTQ raw sequencing reads were mapped to the NCBI Mus musculus reference genome (Build mm39) using the BWA-MEM algorithm (v0.7.17)[16]. The alignment pipeline included marking duplicate reads with Samblaster (v0.1.24)[17], followed by sorting and indexing the resulting BAM files using Samtools (v1.9)[18]. Depth of coverage across the BAM files was estimated using Mosdepth (v0.3.3). Two complementary variant calling methods were employed to ensure robust identification of SNVs and INDELs. The first method used GATK HaplotypeCaller (v4.1.8.1), followed by joint genotyping with GenotypeGVCF[19]. The second approach relied on Strelka2 (v2.9.10)[20] configured for germline variant calling. To maximize specificity, only variants marked as “PASS” by both tools were retained, and overlapping variants between the two sets were identified using BCFtools (v1.3.1)[21], and any variants present in the mm39 blacklist were excluded. The functional impact of genetic variants was annotated using the VariantAnnotation R package (v3.20)[22]. Variant calls in VCF format were analysed against the TxDb.Mmusculus.UCSC.mm39.knownGene transcript database and the BSgenome.Mmusculus.UCSC.mm39 reference genome.

### Mutational Signature Analysis

Mutational signatures for SBS and ID were extracted using SigProfilerExtractor (v1.1.4)[23], a tool that applies non-negative matrix factorization (NMF) to determine the optimal number of mutational signatures in a dataset. Signatures were identified de novo and matched to the COSMIC mutational signature database (v3)[24], retaining those with a cosine similarity score above 0.9. All associated code is publicly accessible at https://gitlab.com/pasquali-lab/mouse-wgs.

---

**Supplementary Figure 1.** (A) Western blot showing decreased expression of MLH1 protein in shMLH1^KP^ compared with WT^KP^ cell line. β-actin used as a loading control. (B) Gating strategies used for analysis and cell sorting of cancer cells, tumour-infiltrating T cells and myeloid cells and (C) Eos, eosinophils; TAM, tumour-associated macrophages; cDC, dendritic cells. (D) Percentage of TAM A, TAM B or TAM C cells in the total CD45^+^ cell population, evaluated by cytometry. N=13 for IgG2a and N=16 for anti-PD1 treated mice (E) Percentage of dendritic cells (cDC1 or cDC2) in the total CD45^+^ cell population, evaluated by cytometry. N=8 per cohort.

**Supplementary Figure 2.** (A) Evaluation of immunogenicity of single cell clones derived from WTKP cell line by measuring tumour weight (mg) after subcutaneous implantation in immunocompetent (IC) or immunodeficient (ID) mice. N=3 independent single KP cell clones, N=1 tumour per clone per mice. **P≤ 0.01, two-tailed unpaired t-test. (B) qRT-PCR results showing decreased expression of MLH1 protein in sgMLH1^KP^ compared with WT^KP^ cell line. β-actin used as a housekeeping gene. **P≤ 0.01, two-tailed unpaired t-test. (C) Western blot showing decreased expression of MLH1 protein in sgMLH1^KP^ compared with WT^KP^ cell line. β-actin used as a loading control. (D) Percentage of TAM A, TAM B or TAM C cells in the total CD45+ cell population, evaluated by flow cytometry. N=9 IgG2a, N=7 anti-PD1-treated mice. (E) Percentage of dendritic cells (cDC1 or cDC2) in the total cDC cell population, evaluated by flow cytometry. N=9 IgG2a, N=7 anti-PD1treated mice.

## Disclosure and competing interests

The authors declare that they have no conflict of interest.

## Authors’ contributions

E.A., I.Z., and A.K., Conceptualization. Data curation. Formal analysis. Investigation. Methodology. Validation. Visualization. Project administration. Writing - original draft. Writing - review & editing. D.R. Data curation. Formal analysis. Validation. Visualization. M.S.G. Data curation. Formal analysis. Investigation. Methodology. Software. Validation. Visualization. Writing - original draft. Writing - review & editing. L.P. Conceptualization. Supervision of M.S.G. Writing - review & editing. J.A, C.R.L and A.J. Conceptualization. Visualization. Project administration. Writing - original draft. Writing - review & editing. Supervision. Resources. Funding acquisition.

## Acknowledgements

We thank Dr A. Celià-Terrassa, and Dr. K. Sutherland for providing reagents; Dr. Yacine Kharraz and CRG/UPF Flow Cytometry Unit with sample preparation and analysis. This work was supported by grants and fellowships from the Spanish Ministry of Science and Innovation Grant to A.J. (PID2021-127710OB-I00), “La Caixa” foundation (51110009 and HR22-00402). A.J. is supported by Ramon y Cajal Research Fellowship (RYC2018-025244-I). I.Z. is funded by an AECC Postdoctoral Fellowship (POSTD234858ZADR). C.L.R. and J.A. were funded by grants from the Spanish Ministry of Science and Innovation (PID2021-128721OB-I00), and Worldwide Cancer Research (20-0144). A.K. was supported by a predoctoral fellowship of the Spanish Ministry of Science and Innovation (PRE2019-088166). This work was made possible through the “Unidad de Excelencia María de Maeztu’’ funded by the MCIN and AEI (CEX2018-000792-M).

## Notes

### Competing Interest Statement

The authors have declared no competing interest.

