## Supplementary figures and images for "Site-specific immunogenicity and anti-PD1 response in mismatch repair deficient lung adenocarcinoma models"

### Supplementary Figure 1

Supplementary Figure 1

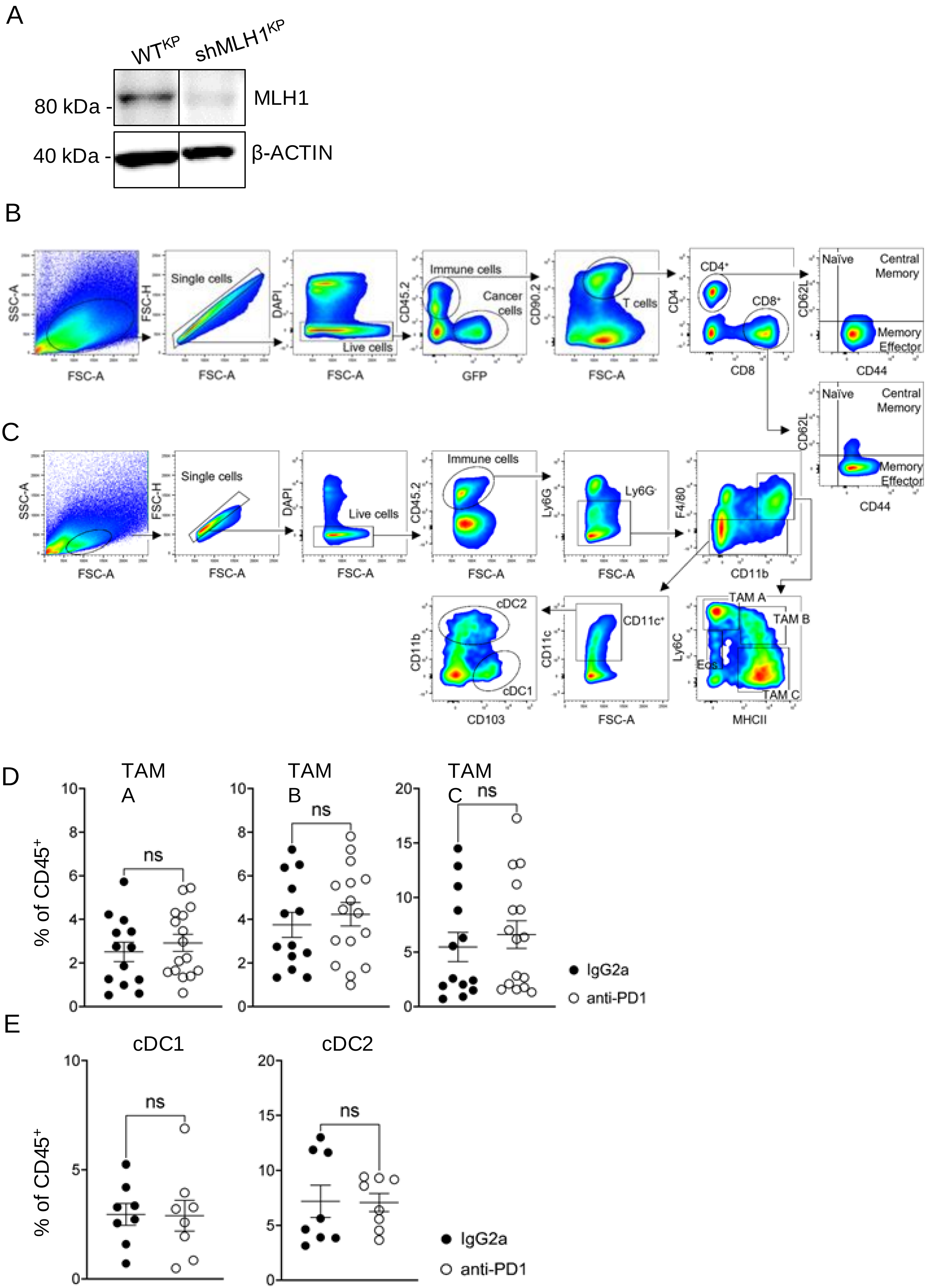

### Supplementary Figure 2

Supplementary Figure 2

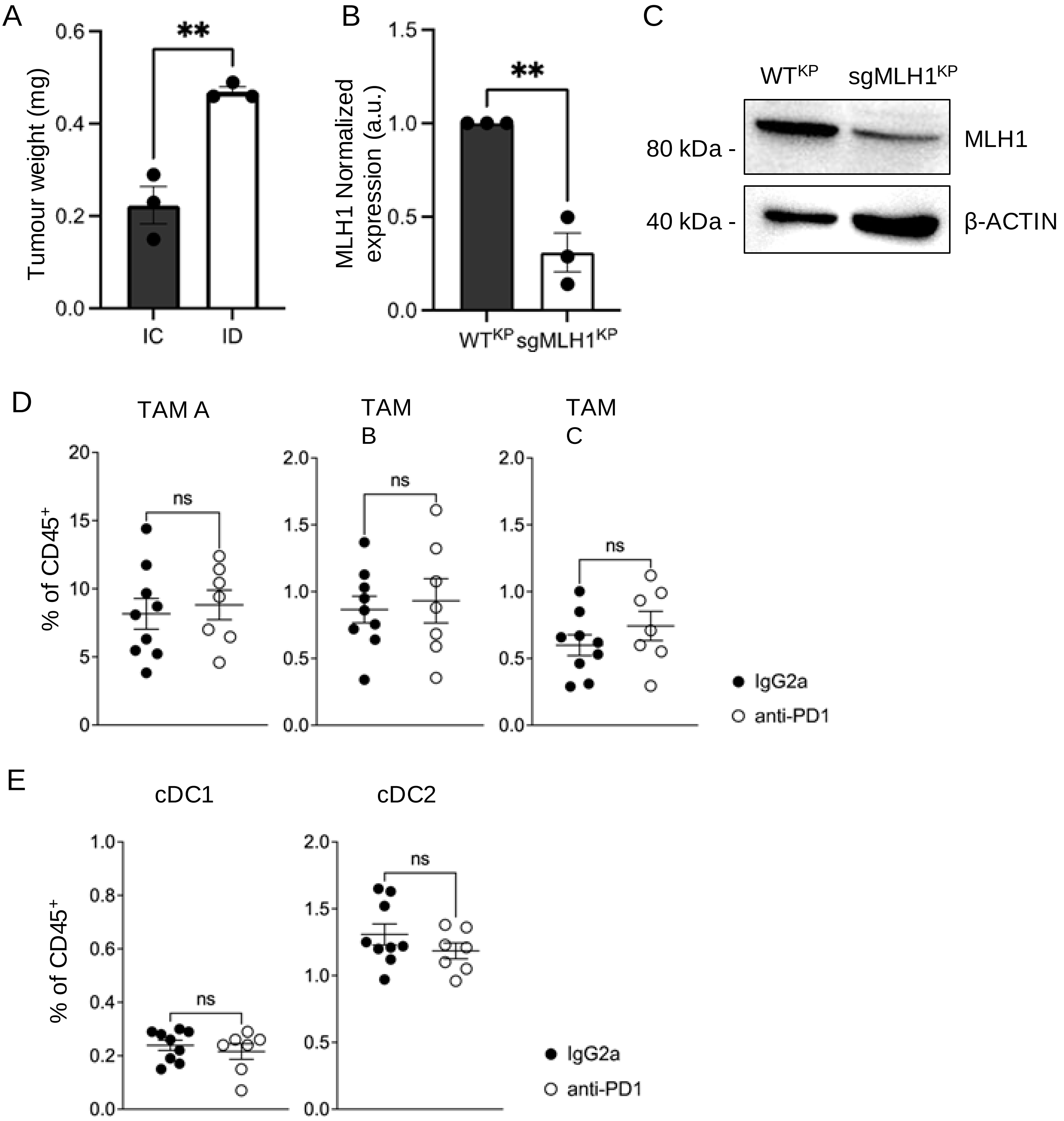
